# Comprehensive Analysis of Exportins Expression and its Prognostic Significance in Colon Adenocarcinoma: Insights from Public Databases

**DOI:** 10.1101/2024.07.18.604060

**Authors:** Punita Kalia, Rohini Ravindran Nair, Suresh Singh Yadav

## Abstract

Colon cancer remains a significant health burden globally, necessitating deeper investigation. Identification and targeting of prognostic markers can significantly improve the current therapeutic approaches for colon cancer. The differential nuclear transport (import and export) of cellular proteins, plays an important role in tumor progression. Exportins, critical mediators of nuclear export, have emerged as potential players in cancer pathogenesis. However, their precise roles and prognostic significance in colon adenocarcinoma remain elusive. This study was designed to comprehensively analyse the expression and prognostic significance of all seven exportins in Colon Adenocarcinoma (COAD) using the online public database. We used public databases UALCAN, C-Bio portal, Human Protein Atlas (HPA), and DAVID, to investigate exportins in COAD patients. Kaplan-Meier plotter, Gene ontology (GO), TIMER, STRING, and KEGG were used to analyse data and draw conclusions. Our observations showed a significant correlation of exportins expression with clinical parameters, used to predict a patient’s prognosis in general, such as advancing tumor stage, overall/relapse-free survival, and immune cell infiltrations. Mutation analysis showed the presence of amplifications, missense mutations in XPO2 and XPO4, and deep deletions in XPO7 genes contributing to disease progression and patients survival. This study highlights the potential use of exportins as novel prognostic biomarkers and therapeutic targets for colon adenocarcinoma progression and management.

## INTRODUCTION

The colon and rectum are parts of the large intestine, and their cancers are known as colon cancer and rectal cancer, respectively. These cancers are often collectively referred to as colorectal cancer. Colorectal cancer ranks among the most prevalent cancers worldwide, standing as the third most diagnosed globally and second leading cause of cancer-related deaths globally, contributing to 10% of all cancer cases, as per the latest WHO report. The predominant type of colorectal cancer is colorectal adenocarcinoma which usually emerges from the glandular epithelial cells of the large intestine, accounting for 95% of all cases [1][2]. Similarly, the predominant type of colon cancer is Colon adenocarcinoma.

Cellular protein functions in multiple locations inside the cells which is made feasible with the help of transporter proteins. Improper localization of proteins either leads to their inactivation or over-activation, which can hamper cellular functioning responsible for ensuring their correct positioning within the cell. A well-identified transporter is karyopherin which is involved in nucleocytoplasmic shuttling of proteins. Karyopharin can be categorized as exportins and importins. Importins facilitate the entry of cargo proteins into the nucleus while exportins mediate the exit of cargo proteins and RNA, such as mRNA, tRNA, and rRNA, from the nucleus. Cargo proteins, to be exported from the nucleus, interact with exportin through a typically leucine-rich nuclear export signal (NES) [3]. Karyopherin expression has been analysed in different types of cancer [4]. In the current study, we have focused on the role of exportin and its association with colon adenocarcinoma. Additionally, these proteins also play crucial roles in mitosis and transcription. Therefore, it is important to analyse the differential expression levels of exportins in diseases, regardless of their function as transporter or non-transporter.

Major exportins studied to date are XPO1 (CRM1), XPO2 (CSE1L), XPO5, and XPOT however, very limited information is available on XPO4, XPO6, and XPO7 in cancer progression. XPO1 has been implicated in the progression of several cancer types, including pancreatic, glioma, lung, ovarian, gastric, colorectal, kidney, and prostate cancer [5–12]. XPO2 is found to be significantly overexpressed in a variety of cancers including oral cancer, gastric cancer, lung cancer, colorectal cancer, laryngeal cancer, breast cancer, osteosarcoma, chronic myeloid leukaemia, ovarian cancer, hepatocellular carcinoma [8,13–20]. XPO5 dysregulation is also observed in colorectal cancer, breast cancer, non-small cell lung carcinoma, prostate cancer, and thyroid cancer [13,21–24]. XPOT is upregulated in hepatocellular carcinoma, neuroblastoma, and triple-negative breast cancer [25–27]. XPO4 is upregulated in prostate cancer and downregulated in hepatocellular carcinoma [28,29]. XPO6 has been exclusively identified as an oncogene in prostate and breast cancer [30,31]. XPO7 was also found to be dysregulated in ovarian cancer, prostate cancer, liver cancer, and breast cancer [27,32–34]. Inhibitors of nucleocytoplasmic transport are continuously being investigated in cancer [35][36]. The most noticeable is LMB (Leptomycin B) which inhibits the interaction of XPO1 with its cargo protein. LMB induces apoptosis in cancer cell lines, but cell cycle arrest in normal lung fibroblast [37]. Seervi et al. demonstrated that inhibiting XPO1 significantly reduced the recovery of apoptotic cells in anastatic cancer cells [38]. Anastatic cancer cells are cells that resist apoptosis even after the strong apoptotic signal. This suggests the significance of these transporter proteins in cancer therapeutics. However, it is poorly explored in colon cancer progression and management.

To our knowledge, no studies have addressed exportin expression as a prognostic marker in colorectal cancer or colon adenocarcinoma, except for one reported by Wang et al. in 2022. Wang et al. demonstrated that high expression of XPO2 (CSE1L) was a risk factor for colorectal cancer patients with poor prognosis [39]. Karyopherin has been shown to significantly affect the prognosis of patients with malignant germ cell tumors [5]. The present study has comprehensively addressed the expression pattern and prognostic significance of all major exportin (XPO1, XPO2, XPO4, XPO5, XPO6, XPO7 and XPOT) in colon adenocarcinoma. Our analysis includes expression profiling, genetic alterations, and, immune infiltration to evaluate the prognostic significance of exportins in colon adenocarcinoma. We also provide the protein-protein interaction network suggesting exportin’s co-co-expressed, co-regulated, and functionally related proteins. Our observations suggest that exportins are differentially expressed, have frequent genetic alterations, and have several functions as oncogenes in colon adenocarcinoma. Their correlations with the patient’s clinical parameters pose them to be potential prognostic factors in colon adenocarcinoma.

## METHODOLOGY

### Data Collection

We used the COAD dataset for expression at the mRNA level and protein level from The Cancer Genome Atlas (TCGA) and Human Protein Atlas (HPA) databases respectively. A total of 45 normal and 274 tumor patient samples from the TCGA database were utilised in ULCAN analysis. A total of 1624 patients’ data for overall survival and 1148 patients’ data for relapse-free survival was used in KAPLAN-MEIER PLOTTER analysis from multiple databases including GEO, EGA, and TGCA (supplementary table 1). TCGA Firehose Legacy module of the cBioPortal database was used for genetic alteration analysis. It contains 640 samples (392 colon adenocarcinoma, 169 rectal adenocarcinoma, 66 mucinous colon adenocarcinoma, and 13 mucinous rectal adenocarcinoma samples). Our analysis was restricted to the 392 colon adenocarcinoma samples. The COAD dataset of the TIMER database was used to analyse the correlation between exportin expression and tumor immune cell infiltration in colon adenocarcinoma cancer.

### UALCAN analysis

This interactive web resource (http://ualcan.path.uab.edu/) enables the analysis of cancer omics data. It primarily uses the data from The Cancer Genome Atlas (TCGA) database. TCGA provides clinical patient data and allows to analyse of expression, survival, methylation, correlation, and pan-cancer view of 33 different cancer datasets [40] Gene expression levels were quantified as transcripts per million (TPM), providing a relative measure that facilitates comparison across samples. Statistical significance was determined at a threshold of p<0.05.

### KAPLAN-MEIER PLOTTER analysis

It is an online tool (https://kmplot.com/analysis/) used for patient survival analysis, primarily focusing on gene expression data to assess the impact of specific genes on the survival of patients. It integrates data from multiple databases, including GEO, EGA, and TGCA [41]. We utilized this to examine the prognostic relevance of Overall Survival (OS) and Relapse-Free Survival (RFS) by analysing the exportins mRNA expression database. Evaluation of Hazard ratio with a confidence interval of 95% and log-rank P value was used to present overall survival/ and relapse-free survival between patients with low and high exportin expression. Differences were considered statistically significant with P<0.05.

### HUMAN PROTEIN ATLAS (HPA) analysis

It is an online database (https://www.proteinatlas.org/) to map all human proteins in cells, tissue, and organs using omics technology, antibody-based imaging, mass spectrometry-based imaging, and transcriptomics [42] Using the pathology section of HPA we compared the protein expression of exportins between normal and colon adenocarcinoma patients.

### cBioPortal for Cancer Genomics

It is a web-based resource (https://www.cbioportal.org/) that provides visualization, analysis, and exploration tools for multidimensional cancer genomics data sets. It allows us to interactively explore genetic alterations, and other molecular characteristics across various cancer types and patient cohorts [43] Here in this study we analysed the mutations of 7 different exportins in the COAD dataset.

### TIMER (tumor immune estimation resource) analysis

TIMER (http://timer.comp-genomics.org/timer/) is a comprehensive online resource for systemically analysing immune cell infiltrates in several cancer types. It is a powerful tool for the investigation of tumor-immune interactions, utilizing data from The Cancer Genome Atlas (TCGA) [44]. Using TIMER we analysed the correlation of exportins expression in tumors with the level of different immune cell infiltration. Statistical significance was determined at a threshold of p<0.05.

### STRING analysis

The STRING database (https://string-db.org/) is a comprehensive online resource that provides information about protein-protein interactions. It aims to collect, store, and integrate available sources including experimental data, computational predictions, and publicly available databases [45] The ten most highly co-expressed genes for each exportin were collected from the co-expression module in the cBioPortal database. Then these gewere were used for STRING analysis to draw the network of protein-protein interaction. Spearman’s correlation was used with P < 0.001.

### DAVID database for gene ontology and KEGG pathway analysis

It is an online database (https://david.ncifcrf.gov/summary.jsp) for functional annotations and enrichment analysis [46]. We performed gene ontology (GO) and KEGG pathway analysis of exportins and their 70 co-expressed genes in DAVID. Statistical significance was determined at a threshold of p<0.05.

## RESULTS

### Exportins expression in colon adenocarcinoma patients

To explore the exportins expression in colon adenocarcinoma patients, we conducted UALCAN analysis on seven exportins (XPO1, XPO2, XPO4, XPO5, XPO6, XPO7, XPOT). We observed overexpression of all the exportins analysed in colon adenocarcinoma patients compared to normal individuals (Figure 1A-H). We also analysed the differential expression of all exportins using the TNMplot database and observed similar results (supplementary figures 1 and 2). Protein expression of various exportins was analysed using the Human Protein Atlas (HPA) database. We compared the protein expression of exportins in COAD tissue samples with normal human colon tissue samples. Moderate staining for XPO2, XPO5, and XPO6 was observed in COAD samples, while normal samples exhibited low or undetectable staining. In COAD samples, low staining of XPO7 was observed, whereas normal tissues showed no staining. Conversely, XPO1 exhibited high staining in normal samples compared to COAD samples (Figure 2). Protein expression data for XPO4 and XPOT were not available in the HPA. Overall, this expression analysis reveals an upregulation of exportin proteins in COAD, except for XPO1, as illustrated in Figure 2.

**Figure 1.**
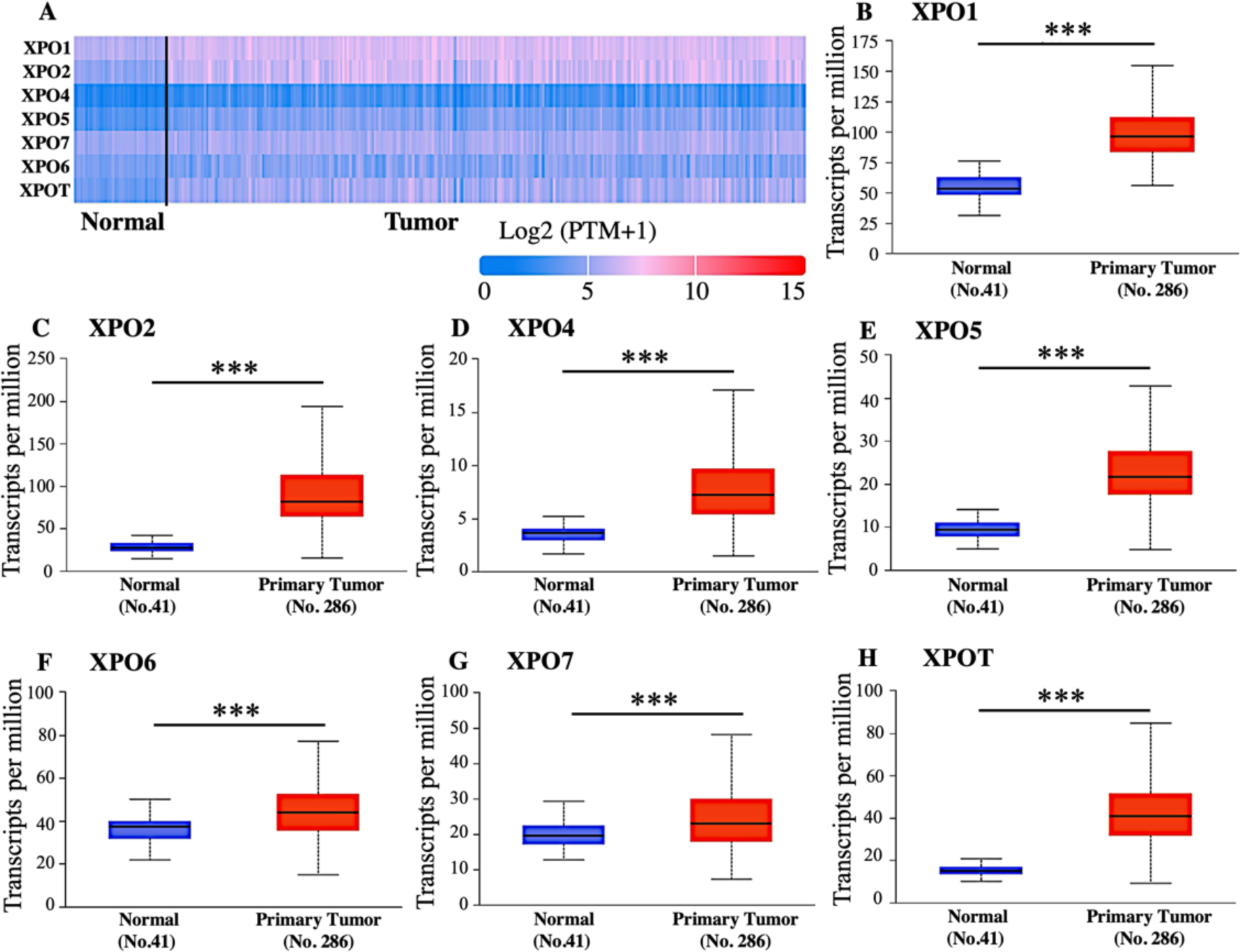
**A.** Differential mRNA Expression of exportins in Colon Adenocarcinoma using TCGA database: A: Heatmap from UALCAN database shows the mRNA expression of exportins. B-H. The box plot illustrates the mRNA expression levels of exportins (XPO1, XPO2, XPO4, XPO5, XPO6, XPO7, and XPOT) in colon adenocarcinoma samples (Tumor) compared to normal (N) colon tissue samples. Expression level is represented as transcripts per million. Significant differences in expression between tumor and normal tissues are indicated by asterisks (*p < 0.05, **p < 0.01, ***p < 0.001).

**Figure 2:**
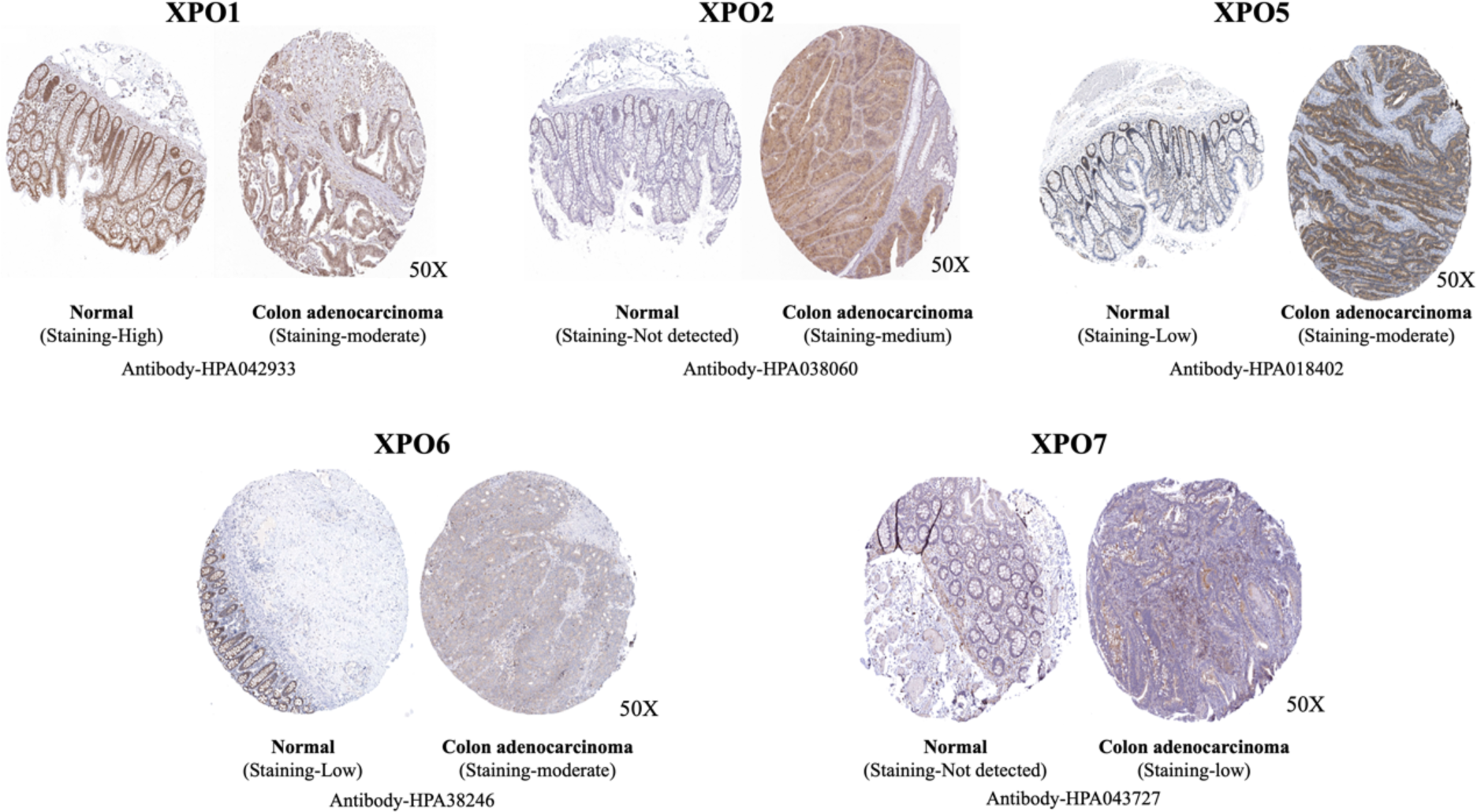
Immunohistochemistry analysis of exportins in Colon Adenocarcinoma using HPA (Human Protein Atlas) database: Representative picture showing expression of XPO1, XPO2, XPO5, XPO6, XPO7 in Colon Adenocarcinoma and control tissue.

### Exportins expression and its Association with tumor stage and patient survival

We conducted UALCAN analysis to demonstrate the correlation of clinical parameters such as tumor stages and patient survival with the expression of exportins in COAD patients to understand the role of exportins in the prognosis of COAD patients. As shown in Figure 3, we observed that stage 4 tumors show the highest median expression levels of exportins XPO1, XPO4, XPO5, and XPOT, suggesting that their expression increases with cancer progression. For XPO4, expression levels are significantly higher in stage 4 compared to stage 2, while XPO5 shows significantly higher expression in stage 3 than in stage 2 (Figure 3). Notably, XPO7 exhibits a consistent and significant decrease in expression with advancing tumor stage, suggesting a negative correlation between tumor stage and its expression levels in COAD patients. The expression of XPO2, XPO6, and XPOT did not exhibit a conclusive trend with increasing tumor stage.

**Figure 3:**
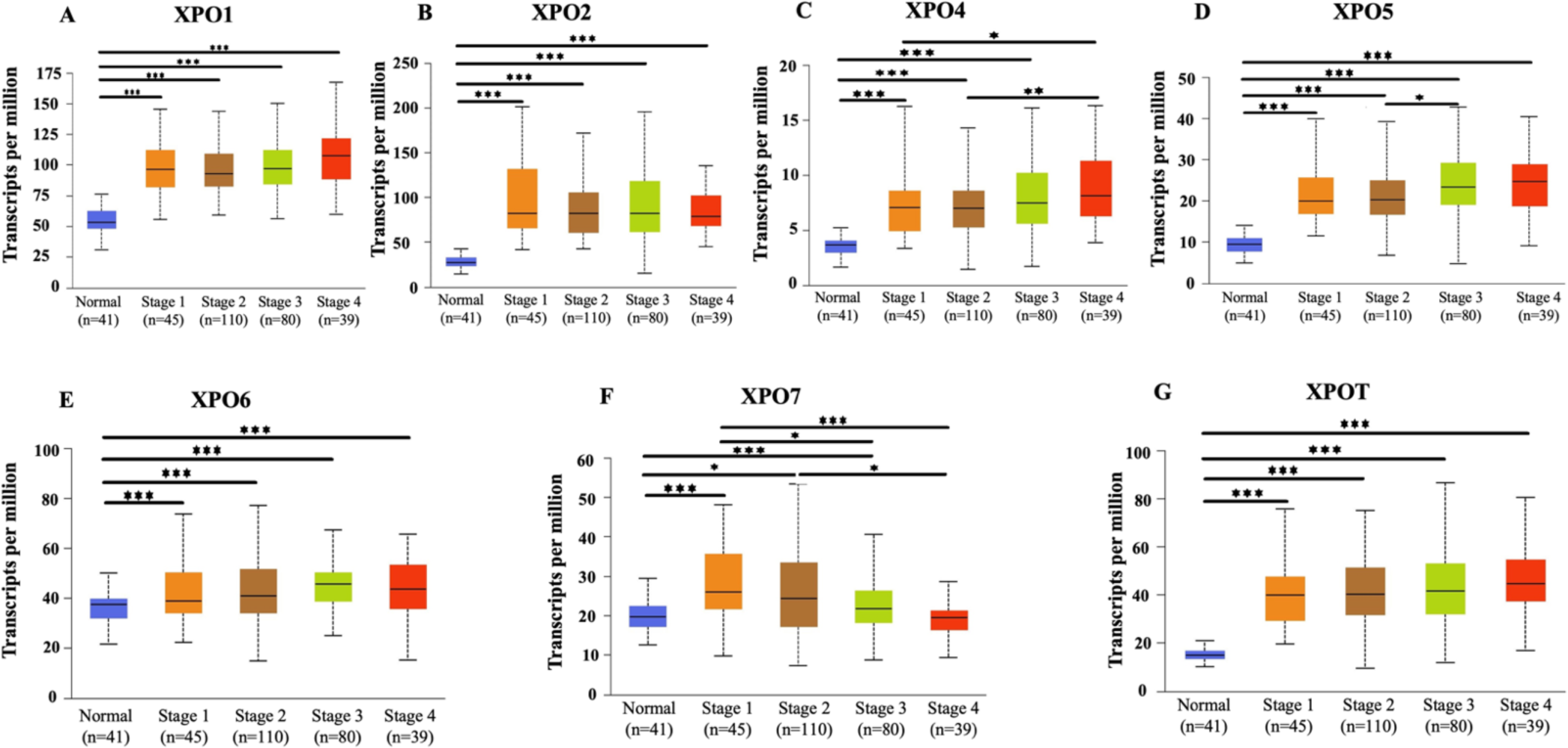
Box plot showing a correlation between exportins (XPO1, XPO2, XPO4, XPO5, XPO6, XPO7, and XPOT) mRNA expression and stage of tumor in Colon Adenocarcinoma using TGCA database in ULCAN analysis. P<0.05*, P<0.01**, P<0.001***.

Further, we divided cancer patients into two groups (high expression level and low expression level of particular exportin based on their median expression level) and performed a survival analysis using the Kaplan-Meier Plotter. We analysed the relationship between mRNA expression of exportins and both overall survival and relapse-free survival of COAD patients. Among the exportins analysed, high expression levels of XPO1, XPO2, XPO4, and XPOT were significantly and negatively associated with the overall survival of the patients. High expression of XPO5 was also negatively associated with overall survival but differences were not significant (Figure 4; Upper panel). Contrary to this the high expression of XPO6 and XPO7 was positively associated with the overall survival of the patients, although the differences were not significant for XPO7. On the other hand, among the exportin analysed expression levels of XPO1, XPO2, XPO4, XPO5, and XPOT were negatively associated with the relapse-free survival of the patients but the differences were significant only for the XPO1 and XPO2. Contrary to this the high expression of XPO6 and XPO7 was positively associated with the relapse-free survival of the patients, although the differences were marginally significant for XPO7 and not significant for XPO6 (Figure 4; Lower panel).

**Figure 4:**
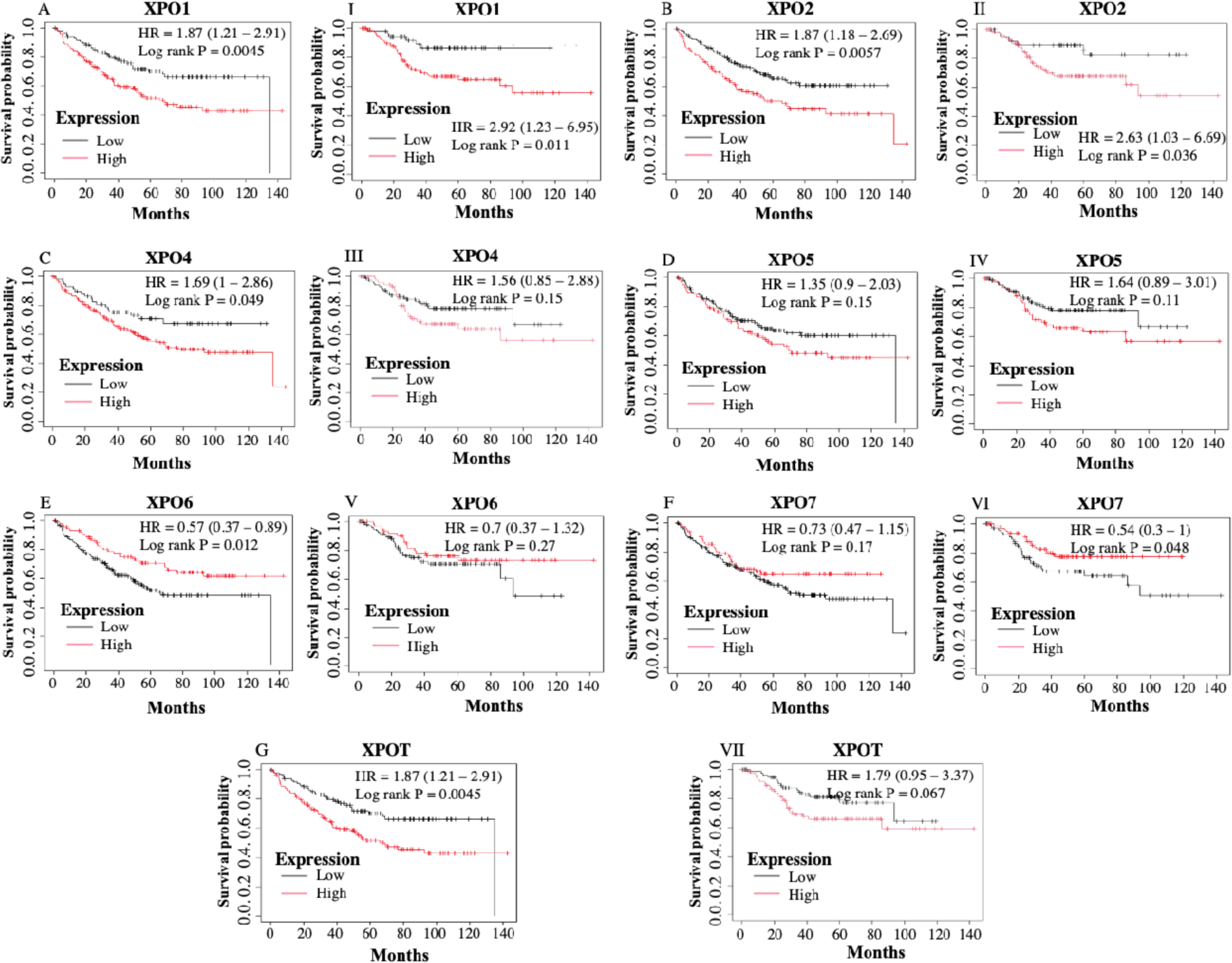
Overall Survival (OS) and Relapse Free Survival (RFS) analysis using Kaplan-Meier Plotter Database in Colon Adenocarcinoma patients: Figure shows the overall survival (A-G) and relapse-free survival (I-VII) of patients with high and low expression levels. Compared to the median expression value, of various exportins (XPO1, XPO2, XPO4, XPO5, XPO6, XPO7, and XPOT). A P-value of <0.05 has been considered as statistically significant.

**Table: 1.** Hazard Ratios (HR) with 95% Confidence Intervals (CI) for overall survival and relapse-free survival of Colon Adenocarcinoma patients with high and low expression levels of exportins: The HR values indicate the relative risk of adverse outcomes associated with the expression levels (high or low compared to their median expression level of respective exportin) of each exportin. An HR greater than 1 suggests a higher risk, while a HR less than 1 suggests a lower risk. The 95% CI provides a range within which the true HR is expected to fall, with 95% confidence.

| EXPORTINS | OVERALL SURVIVAL |  |  | RELAPSE FREE SURVIVAL |  |  |
| --- | --- | --- | --- | --- | --- | --- |
|  | CASES | HR (95% confidence interval) | P value | CASES | HR (95%confidence interval) | P value |
| XPO1 | 232 | 1.78 (1.18-2.69) | 0.0052 | 164 | 2.92 (1.23-6.95) | 0.011 |
| CSE1L | 232 | 1.69 (1-2.86) | 0.049 | 164 | 2.63 (1.03-6.69) | 0.036 |
| XPO4 | 232 | 1.49 (0.97-2.31) | 0.07 | 164 | 1.56 (0.85-2.88) | 0.15 |
| XPO5 | 232 | 1.35 (0.9-2.03) | 0.15 | 164 | 1.64 (0.89-3.01) | 0.11 |
| XPO6 | 232 | 0.57 (0.37-0.89) | 0.012 | 164 | 0.7 (0.37-1.32) | 0.27 |
| XPO7 | 232 | 0.73 (0.47-1.15) | 0.17 | 164 | 0.54 (0.3-1) | 0.048 |
| XPOT | 232 | 1.87 (1.21-2.91) | 0.0045 | 164 | 1.79 (0.95-3.37) | 0.067 |

### Genetic alterations in exportins gene

In the cBioPortal database, COAD patient samples with the TCGA Firehose Legacy module were selected, and the genetic alterations and mRNA microarray data of all exportins were analysed. As shown in Figure 5A, the total genetic alterations rates, out of 634 cases, in each exportin were observed as follows: XPO1: 3% (11 cases), XPO2: 10% (39 cases), XPO4: 8% (31 cases), XPO5: 3% (11 cases), XPO6: 4% (16 cases), XPO7: 8% (31 cases), XPOT: 3% (11 cases). These genetic alterations are missense mutations, truncating mutations, amplifications, and deep deletions, as illustrated in Figure 5A. Figure 5B shows the overall survival of patients with these genetic alterations compared to control. The overall survival of patients with genetic alterations is significantly lower than that of the control group (P=0.03). Figure 5C depicts the contribution of exportins (XPO1, XPO2, XPO4, XPO5, XPO6, XPO7, and XPOT) to various genetic alterations and mRNA expression levels in COAD patients. These alterations levels are as follows: Mutations: 2.37% (9 cases), Amplifications: 9.15% (35 cases), Deep deletions: 3.63% (14 cases), mRNA upregulation: 3.47% (13 cases), mRNA downregulation: 4.73% (18 cases), Multiple alterations: 5.05% (19 cases). The major genetic alteration is amplification, particularly in XPO2, XPO4, and XPO5 (Figure 5A). Deep deletions, which are associated with low mRNA expression, are most commonly seen in XPO7.

**Figure 5:**
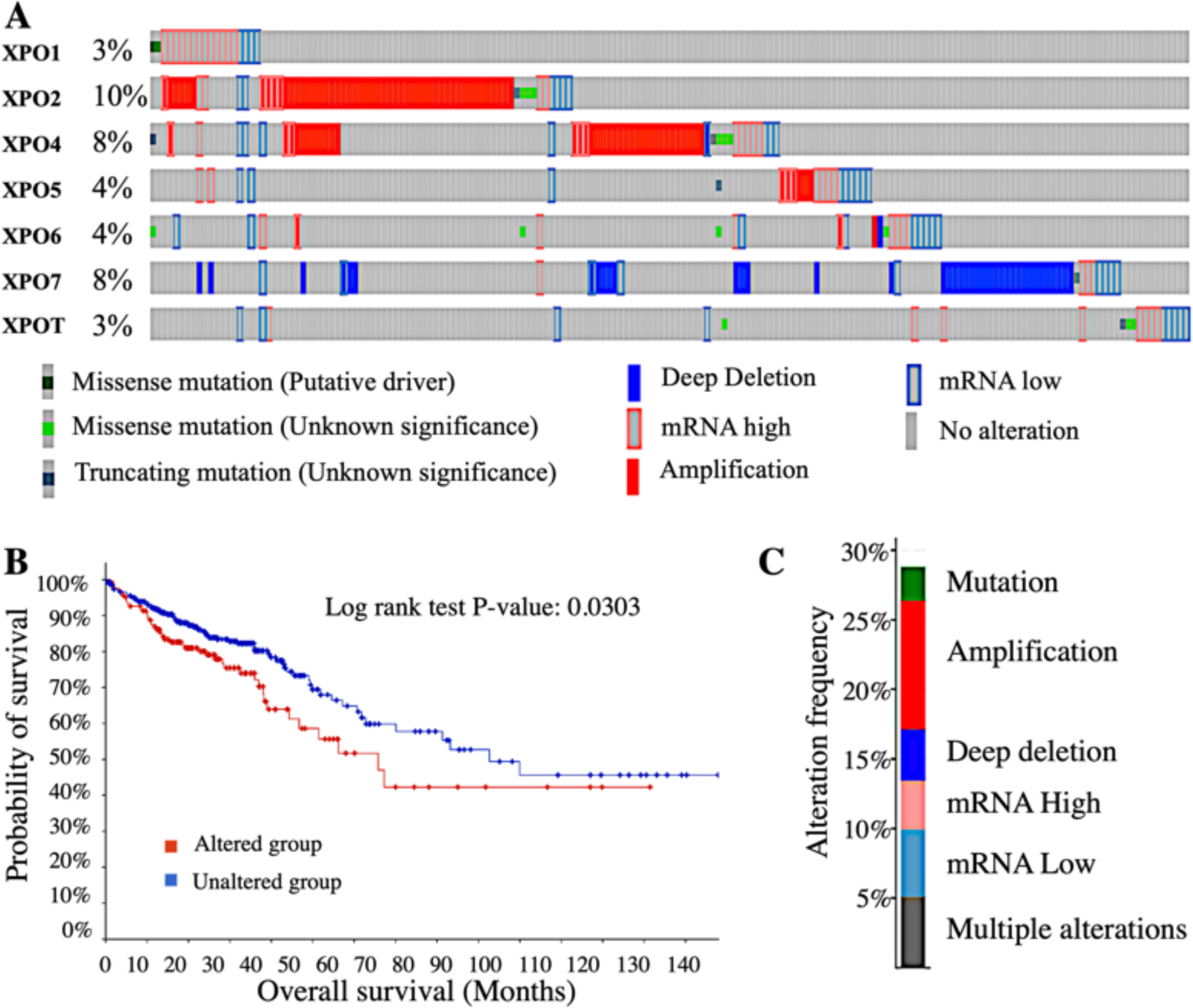
Genetic alteration analysis using cBioportal database in Colon Adenocarcinoma patients. A. showing cBioportal onco-print with percentage of **Genetic alteration in** Colon Adenocarcinoma patient. B. Survival analysis indicates that genetic alterations in exportins are associated with poor overall survival in patients. C. Illustrating the contribution of exportins (XPO1, XPO2, XPO4, XPO5, XPO6, XPO7, and XPOT) to various genetic alterations and mRNA expression levels.

### Correlation of exportins expression with the level of immune cell infiltration in tumor

As we know, the tumor microenvironment plays a major role in cancer progression by providing a supportive environment for tumor growth. Consequently, immune cells in the tumor microenvironment are often suppressed or become tumor-associated. Bu et. al identified four tumor microenvironment-based rectal cancer subtypes and suggested a degenerate gene signature to predict the prognosis and immunotherapy response [47][48]. It has been reported well that the level of infiltrated immune cells in the tumor microenvironment is indicative in predicting the prognosis of patients, thus we investigated it in this study [49] [50].To investigate this, we examined the association of exportins expression with the level of immune cell infiltration in COAD patients’ tumor microenvironment using the TIMER database. As shown in Figure 6: (1) **XPO1:** The only statistically significant correlation observed is between its mRNA expression and tumor purity, with a significant and positive correlation (Rho = 0.12, p = 0.0153). “tumor purity” refers to the proportion of cancer cells in a tumor sample relative to the total number of cells. There are no significant correlations between XPO1 expression and the infiltration levels of the different immune cells analysed such as CD8+ T cells (Rho = -0.016, p = 0.793), CD4+ T cells (Rho = 0.056, p = 0.357), B cells (Rho = - 0.094, p = 0.120), macrophages (Rho = -0.074, p = 0.224), and neutrophils (Rho = -0.047, p = 0.436). Note that, although not significant, XPO2 is negatively correlated with the infiltration of CD8+ T cells, macrophages, and neutrophils. (2) **XPO2:** There is a statistically significant positive correlation between XPO2 expression and tumor purity (Rho = 0.143, p = 0.00392). A statistically significant negative correlation is observed with CD8+ T cell infiltration (Rho = -0.17, p = 0.00458), and neutrophil infiltration (Rho = -0.218, p = 0.000379). Correlation between XPO2 expression and infiltration of CD4+ T cell infiltration (Rho = 0.03, p = 0.619), B-cell (Rho = -0.027, p = 0.658), and macrophage (Rho = 0.043, p = 0.482) was not found to be statistically significant. (3) **XPO4:** There is a statistically significant positive correlation between XPO4 expression and tumor purity (Rho = 0.105, p = 0.0344), while neutrophil infiltration (Rho = 0.043, p = 0.475) showed significant negative correlation. Correlation between XPO4 expression and infiltration of CD8+ T (Rho = -0.031, p = 0.605), CD4+ T cell (Rho = 0.03, p = 0.619), B-cell (Rho = -0.027, p = 0.658) and macrophage (Rho = 0.043, p = 0.482) was not found to be statistically significant. (4) **XPO5:** A statistically significant positive correlation is observed between XPO5 expression and CD4+ T cell infiltration (Rho = 0.319, p = 6.47e-08), and macrophage infiltration (Rho = 0.189, p = 0.0069). However, no significant correlation is found with tumor purity (Rho = 0.076, p = 0.126), CD8+ T cell infiltration (Rho = - 0.078, p = 0.200), B cell infiltration (Rho = -0.007, p = 0.907), and neutrophil infiltration (Rho = 0.05, p = 0.413). (5**) XPO6:** There is a statistically significant positive correlation between XPO6 expression with infiltration of CD8+ T cell (Rho = 0.144, p = 0.0169), CD4+ T cell (Rho = 0.240, p = 0.0000366), macrophage (Rho = 0.223, p = 0.000191), and neutrophil (Rho = 0.287, p = 1.29e-06). However, no significant correlation is found between tumor purity (Rho = 0.01, p = 0.837) and B cell infiltration (Rho = 0.029, p = 0.626). (6) **XPO7:** There is a statistically significant positive correlation between XPO7 expression with infiltration level of CD8+ T-cell (Rho = 0.196; p = 0.00106), CD4+ T-cell (Rho = 0.124, p = 0.0396), Neutrophil (Rho = 0.344, p = 2.50e-09). However, there is no significant correlation with tumor purity, (Rho = -0.041, p = 0.0413), infiltration level of B cell (Rho = 0.036, p= 0.549), and Macrophage (Rho = 0.092, p = 0.126). (7) **XPOT:** There is a statistically significant positive correlation between XPOT expression with infiltration level of CD4+ T-cell (Rho = 0.105, p = 0.0813), macrophage (Rho = 0.175; p = 0.00353) and Neutrophil (Rho = 0.122, p = 0.0428). However, there is no significant correlation with tumor purity, and infiltration level of CD8+ T-cell (Rho = -0.003; p = 0.0964), and B-cells (Rho = -0.055; p = 0.360) (Figure 6).

**Figure 6:**
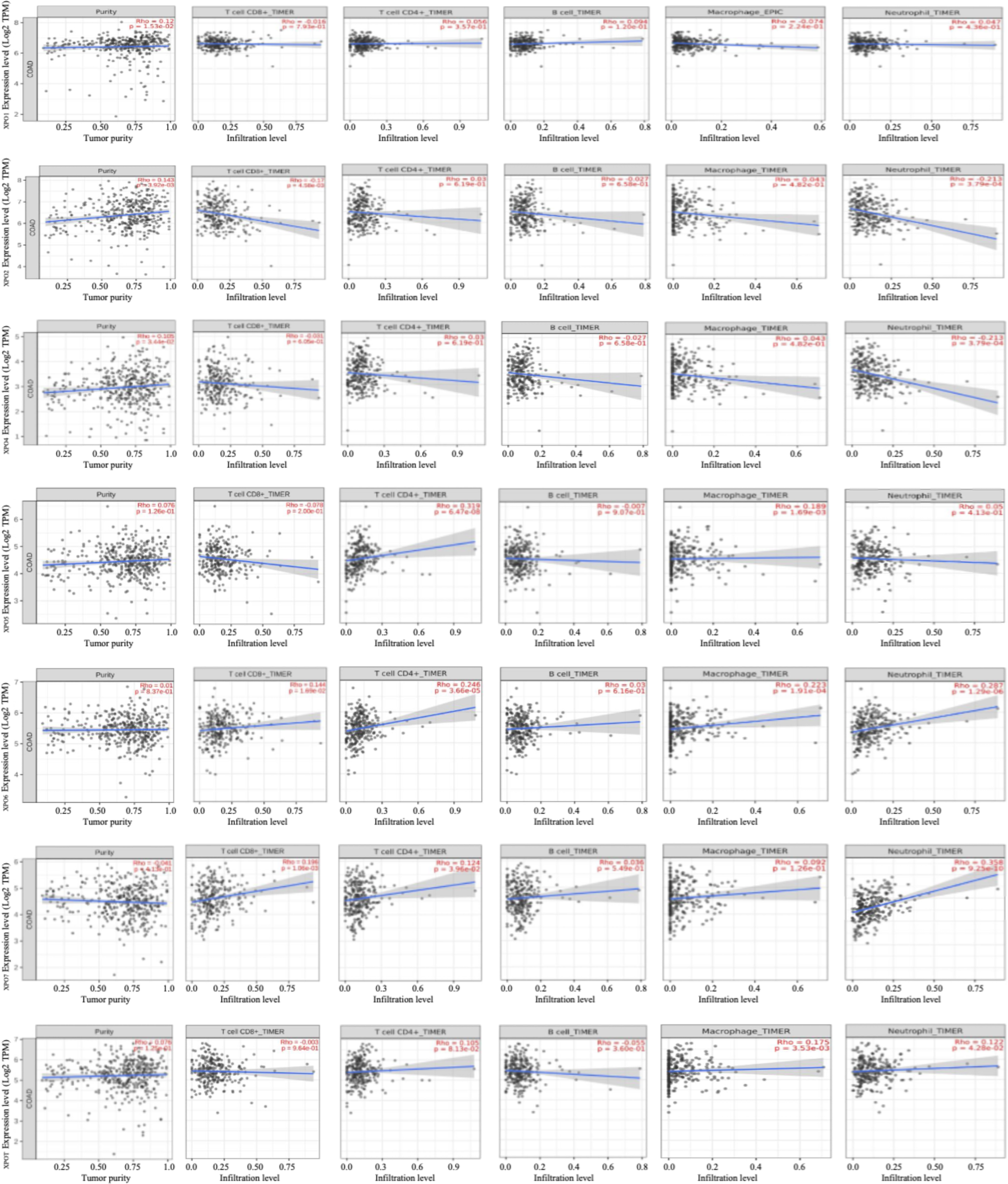
Correlation of exportins expression with the level of infiltrated immune cell using TIMER database: The Y-axis of all the graphs shows the expression (Log2 TPM) of respective exportin and the X-axis shows the level of immune cells (CD8^+^, T-cell; CD4+T-cell; B cell; Macrophage and Neutrophil) infiltration in tumor tissue of Colon Adenocarcinoma patients. The first graph in each panel shows the correlation of exportins expression with the purity of the tumor. “tumor purity” is the proportion of cancer cells out of the total number of cells.

### Co-expressed gene analysis and protein-protein interactions

We used the cBioPortal database to identify the top ten genes that are highly co-expressed with each exportin, using the co-expression module that showed the highest Spearman’s correlation (supplementary table 2). Out of these highly co-expressed genes, the top two co-expressed genes with the highest Spearman’s correlation coefficient for each exportin are as follows: XPO1: ATAD5 and PMS1; XPO2: DPM1 and EIF2S2; XPO4: RNF6 and AKAP11; XPO5: RPL71 and HSP90AB1; XPO6: GTF3C1 and SETD1A; XPO7: CCAR2 and CHAMP7; XPOT: MARS1 and NAA25. Further, using the STRING database and selecting the 10 highly co-expressed genes with the largest Spearman’s rank correlation values, we constructed a protein-protein interaction network for these genes. This analysis makes a map of interactions between exportins and their co-expressed genes (Figure 7 A). Further, using the DAVID database, we tried to categorize at least the top two co-expressed genes of each exportin (a total of 14 genes) based on their involvement in various biological processes. We observed that these genes are primarily involved in processes such as Genome Stability & DNA Repair, Protein Synthesis & Degradation, Signal Transduction & Cell Growth, and Gene Expression & RNA Processing (Figure 7B; Supplementary Table 3). We also prepared a heatmap to show the co-expression of each exportin with the others (supplementary figure 3).

**Figure 7:**
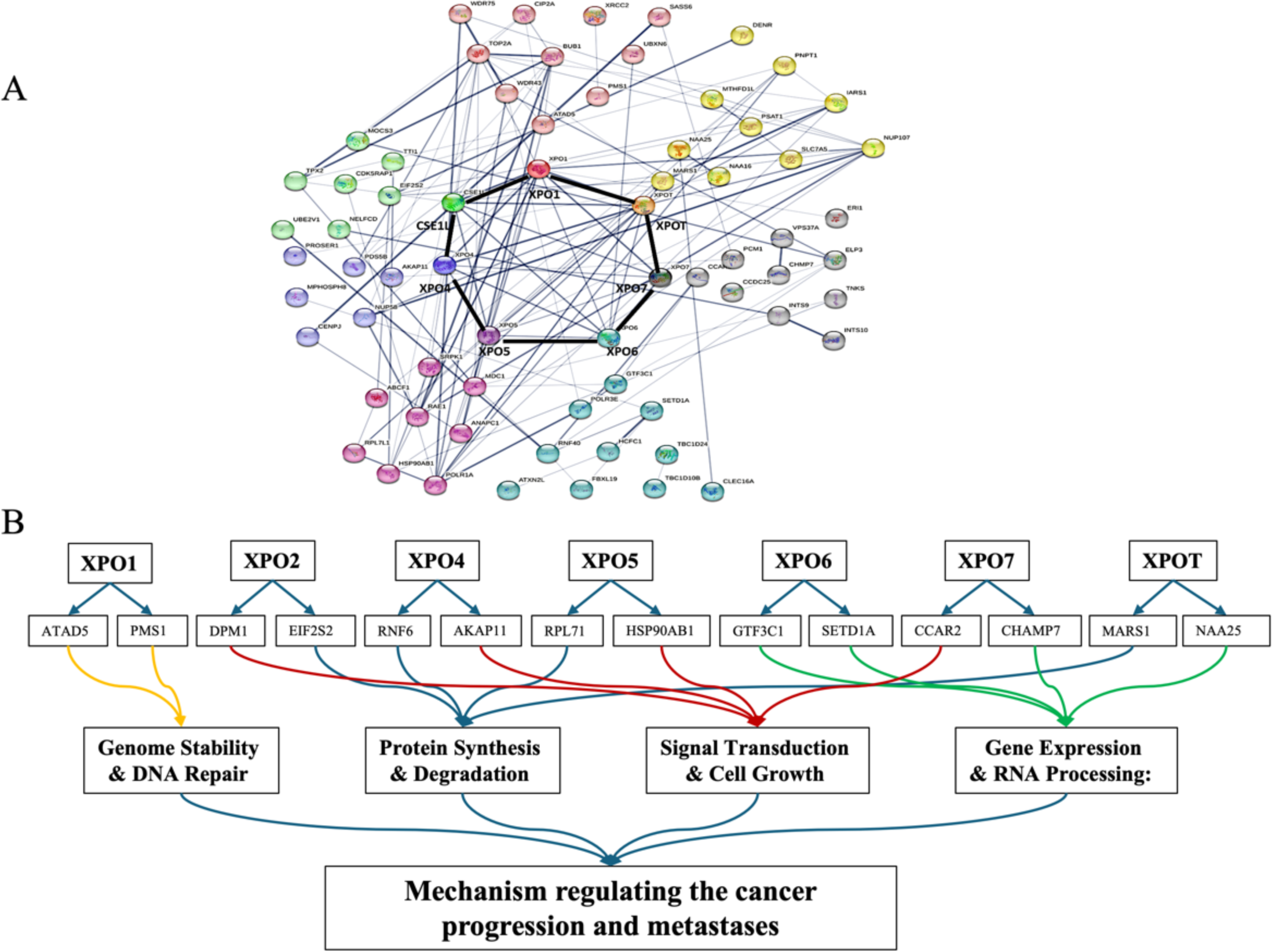
Protein-protein interaction network using STRING analysis: A. Protein-protein interaction network resulting from the total 70 co-expressed genes (10 of each exportin) analysed in STRING. B. Categorising the top two highly co-expressed genes (with the highest Spearman correlation coefficient) to each exportin (total 14) in different biological processes involved in cancer progression.

### Gene ontology and KEGG pathway analysis using the DAVID database

We also performed ontology functional enrichment analysis using the DAVID database, focusing on the top ten biological processes, cellular components, and molecular functions associated with exportins and their co-expressed genes. Processes were considered significant with a p-value of <0.05. As shown in Figure 8A, the significant biological processes mediated by the exportins include: GO:0051301 (Cell cycle), GO:007049 (Cell division), GO:0006974 (Cellular response to DNA damage stimulus), GO:0006338 (Chromatin remodeling), GO:0060236 (Regulation of mitotic spindle organization), GO:0034243 (Regulation of transcription elongation from RNA polymerase II promoter), GO:0045070 (Positive regulation of viral genome replication), GO:0006913 (Nucleocytoplasmic transport), GO:0007059 (Chromosome segregation), and GO:0046601 (Positive regulation of DNA damage checkpoint). Significant cellular components identified include GO:0005829 (Cytosol), GO:0005737 (Cytoplasm), GO:0005654 (Nucleoplasm), GO:0005634 (Nucleus), GO:0005730 (Nucleolus), GO:0005813 (Centrosome), GO:0032991 (Macromolecular complex), GO:0005635 (Nuclear envelope), GO:0005643 (Nuclear pore), and GO:0031965 (Nuclear membrane). Significant molecular functions involving exportins include GO:0005515 (Protein binding), GO:0003723 (RNA binding), GO:0005524 (ATP binding), GO:0045296 (Cadherin binding), GO:0060090 (Binding, bridging), GO:0000175 (3’-5’-exoribonuclease activity), GO:0043022 (Ribosome binding), and GO:0000049 (tRNA binding). Figure 8B represents all gene ontology functions and KEGG pathway analyses. The pathways include hsa03013 (Nucleocytoplasmic transport), hsa04110 (Cell cycle), hsa04914 (Progesterone-mediated oocyte maturation), and hsa05014 (Amyotrophic lateral sclerosis). From these analyses, it is evident that the largest number of genes is associated with the term “Protein binding” under molecular functions (64 genes), indicating a significant role in binding activities. In the cellular component category, “Cytosol” involves the highest number of genes (42 genes), suggesting a high level of cellular activity in the cytosol. Important biological processes that involve exportins in colon adenocarcinoma are Cell division, Cell cycle, DNA damage checkpoints, RNA transport, 3’-5’ exoribonuclease activity, and nucleo-cytoplasmic. It is worth noting that the exportins analysed are also prominently involved in processes that are key targets in present-day cancer therapeutics such as DNA damage checkpoints. (Supplementary table 4)

**Figure 8:**
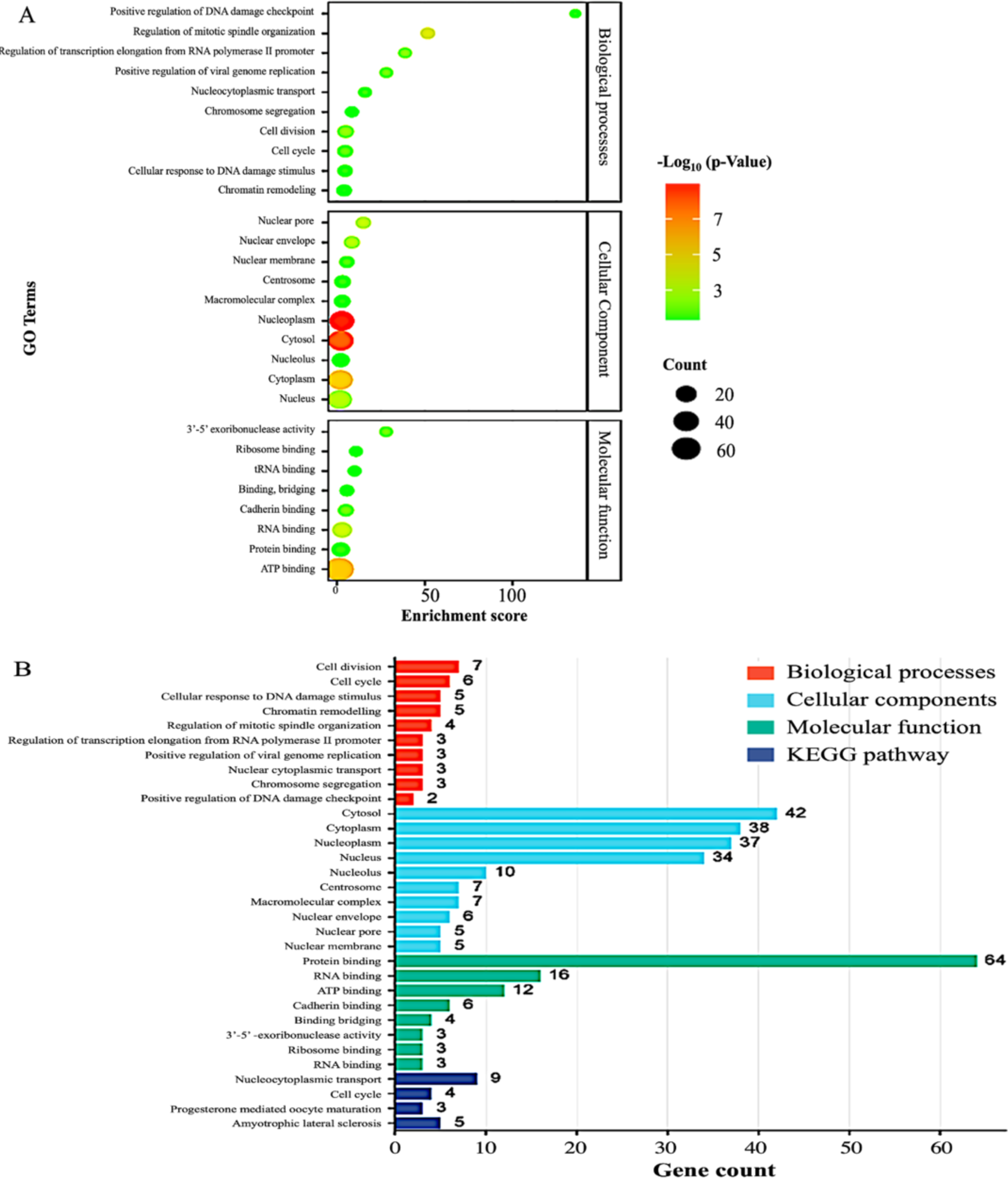
Gene Ontology functional analysis and KEGG pathway analysis: A total of 70 co-expressed genes (10 of each exportin) were analysed using the DAVID database. A. GO enrichment analysis identifying and illustrating the significance of enriched GO terms associated with 70 co-expressed proteins. Each bubble represents a GO term. The size of the bubble corresponds to the number of genes (out of 70) associated with that particular GO term. B. a visual representation of the relationships between GO terms. Gene Ontology graph of biological processes, cellular component, and molecular function, and KEGG pathway analysis.

## DISCUSSION

Exportins are the cellular proteins that primarily facilitate the transport of molecules including oncogenic and tumor suppressor proteins from the nucleus to the cytoplasm and vice versa. Multiple studies have reported the association between exportins expression and tumor progression [51,52]. It has also been reported that depending on the cell and tissue type, exportins can act either as tumor suppressors or tumor promoters [53]. Further investigation is needed to understand the role of exportins in cancer progression and management, especially in Colon Adenocarcinoma. The prognostic values of exportins have been highlighted previously in hepatocellular carcinoma and lung cancer [54] [55]. However, a comprehensive analysis of exportin expression and its prognostic significance in the context of Colon Adenocarcinoma has not been done previously. In this section of the manuscript, we will discuss (1) exportins and their association with tumor stage and patient survival, highlighting their prognostic significance, (2) the prognostic significance of exportins due to their association with level of immune cell infiltration, and (3) protein-protein interaction and exportins mediated pathway analysis.

XPO1 transports various proteins involved in tumor progression, and its expression is significantly higher in cancers [56]. The overexpression of XPO1 mRNA in our analysis is consistent with previous reports. XPO1 overexpression induces the export of p53 out of the nucleus, which facilitates tumor progression and lymph node metastases [11][57]. The prognostic significance of XPO1 protein has been reported in colorectal cancer by Aladhraei et. al [58]. However, our HPA data showed contradictory results for XPO1 expression at the protein level. There is no correlation between mRNA expression and protein levels for XPO1, which may be due to translation regulation or high protein degradation. However, it needs to be validated in further studies. The decreased survival rate in patients with high XPO1 mRNA expression, along with its increased levels in advanced tumor stages, emphasize that Colon Adenocarcinoma patients with high expression of XPO1 would have poor survival. Our observations suggest XPO1 can be used as a potential prognostic biomarker to identify the therapeutic effectiveness of treatment.

Consistent with our observation, XPO2 has been shown to promote tumor growth in colorectal cancer, with its knockdown resulting in reduced cancer progression [20,59]. Another study reports that XPO2 is not only overexpressed in colorectal cancer but also promotes metastasis and invasiveness [60]. Decreased patient survival with high levels of XPO2 highlights its prognostic significance in Colon Adenocarcinoma.

XPO4 is a bidirectional protein transporter capable of importing and exporting cargo protein in and out of the nucleus. In the present study, we found a significant correlation of XPO4 gene expression with tumor stage. Higher expression negatively affects patient survival, indicating its prognostic significance in Colon Adenocarcinoma.

XPO5 is a nuclear export protein responsible for exporting small RNA molecules, particularly miRNA, and double-stranded RNA binding proteins [61]. It has been identified as an oncogene that plays a role in regulating miRNA expression in colorectal cancer [62]. Some reports also suggest it acts as a tumor suppressor, thereby making it an intriguing candidate for cancer therapy [63]. Our observations suggest it functions as an oncogene and has potential prognostic value in Colon Adenocarcinoma, as its expression increases with tumor stage and correlates with poor patient survival.

XPO6 has not been previously studied in colorectal cancer including colon adenocarcinoma, but its prognostic potential has been reported in prostate cancer recurrence [64]. The author showed that relatively high expression of XPO6 is significantly associated with poor patient prognosis. Contrary to this, our analysis reveals its overexpression with better overall survival of colon adenocarcinoma patients. Hence this needs further validation in more samples to better understand it and consider it as a prognostic biomarker.

Our study demonstrated significant overexpression of XPO7 in Colon Adenocarcinoma, suggesting its oncogenic potential. Previous reports have highlighted that XPO7 exhibits dual characteristics: it acts as a tumor suppressor in hepatocellular carcinoma and as an oncogene in prostate cancer. [32,34]. No studies have previously demonstrated the role and expression of XPO7 in Colon Adenocarcinoma. However, its prognostic significance has been reported in serous epithelial ovarian cancer [65]. Our study reveals its overexpression, but decreasing expression with the advancement of the tumor stage, and a significant positive correlation with both overall survival and relapse-free survival, suggesting it is a potential tumor suppressor gene. This pattern suggests three possible roles for XPO7 in Colon Adenocarcinoma: (1) Initiation of Cancer: XPO7 is likely involved primarily in the initial stage to trigger cancer initiation. (2) Tumor Suppressor Function: The decrease in XPO7 expression with tumor progression might be due to its tumor suppressor function, as previously reported [34]; (3) Oncogene-induced senescence: XPO7 probably promotes oncogene-induced cell senescence in cancer cells, leading to tumor suppression and better patient survival. This role has been previously reported by Andrew J. Innes et al [66]. Senescence induction could limit tumor growth and improve clinical outcomes for patients. Andrew J. Innes et. al. showed that XPO7 is a novel regulator of senescence and validated its function in replicative, and oncogene-induced senescence. As reported previously, oncogene-induced senescence primarily acts as a barrier to cancer development by inducing growth arrest in cancer cells, under certain conditions [67]. We speculate similarly in the case of XPO6, as very limited information is available on it. Thus, colon adenocarcinoma patients with high exportin expression are associated with worse overall survival and tumor relapse-free survival, implicating exportins in the aggressiveness and poor prognosis of the disease. XPOT has been identified as a prognostic predictor and potential therapeutic target in neuroblastoma, with its knockdown inhibiting cell proliferation and migration [68]. Similarly, our findings indicate its oncogenic role in colon adenocarcinoma, where its elevated expression is associated with poor prognosis. Our above-mentioned observations validate exportins as potential biomarkers for prognosis and targets for therapeutic intervention in Colon Adenocarcinoma. A high number of genomic alterations in exportins suggests the presence of abnormal exportin protein, and its altered regulation at the genome level, majorly contributing to the cancer progression and patient survival.

The level of immune cell infiltration in the tumor microenvironment is crucial for the better prognosis of the patients [69] [70]. CD8+ T-cells and neutrophils are key immune cells that play a major role in tumor cell death. The blood neutrophil-to-lymphocyte ratio (NLR) serves as a prognostic factor in solid tumors, being negatively associated with patient prognosis [71,72]. Ohashi et al. also demonstrated a negative correlation between NLR and the levels of immune cells infiltrating lung cancer tissues. The presence of higher levels of infiltrating CD8+ T-cells in tumors is generally predictive of a better prognosis, while higher neutrophil levels are indicative of a worse prognosis. Our findings show a significant negative correlation between the expression of exportins (XPO2 and XPO4) and the levels of CD8+ T-cell and neutrophil infiltrations. It suggests that exportins could serve as prognostic markers as it is directly associated with the NLR in tumors. Similar to our previous discussion, the positive association of XPO 6 & 7 expressions with the level of CD8^+^ T-cell infiltrations suggests it is a tumor suppressor in Colon Adenocarcinoma. The association between XPO1, XPO5, and XPOT expression and level of CD8^+^ T-cells infiltration was not significant, so cannot be concluded in this study.

Most of the top co-expressed genes associated with the exportins, as shown in our STRING network analysis, are either directly or indirectly involved in major mechanisms regulating cancer progression and metastases. The co-expressed genes are correlated to other exportins too suggesting the complex functional interactions to regulate cancer progression and metastases. Considering the gene function information in STRING, the top 2 co-expressed genes, according to the Spearman coefficient, can be grouped primarily into four cellular processes as shown in Figure 7B. These processes are predominantly involved in cancer progression. Gene ontology, functional enrichment analysis, and KEGG pathway analysis provide information that exportins analysed are related to the cell cycle, cell proliferation, signal transduction, and repair pathways, in addition to nucleo-cytoplasmic transports.

From our studies, we conclude that exportin expression can be used as a prognostic biomarker for colon adenocarcinoma progression and management. However, there are several open questions to elaborate on the role of exportins in cancer progression and other diseases such as; How genetic alterations affect the binding of cargo protein with exportins?, and the mechanism of XPO7 downregulation at different stages of cancer which alleviates oncogene-induced senescence. Whether it’s a transcription factor too for the genes involved in carcinogenesis. After all our comprehensive analysis opens an avenue to explore several aspects of exportins in cancer research and therapeutics.

## Statements and Declarations

### Author Contributions

PK: Data mining, analysis, and manuscript writing; RRN: Data analysis and revision of the manuscript for important intellectual content; SSY: Study conception and design, data analysis and interpretation, and manuscript writing.

