## Supplementary data for "Comprehensive Analysis of Exportins Expression and its Prognostic Significance in Colon Adenocarcinoma: Insights from Public Databases"

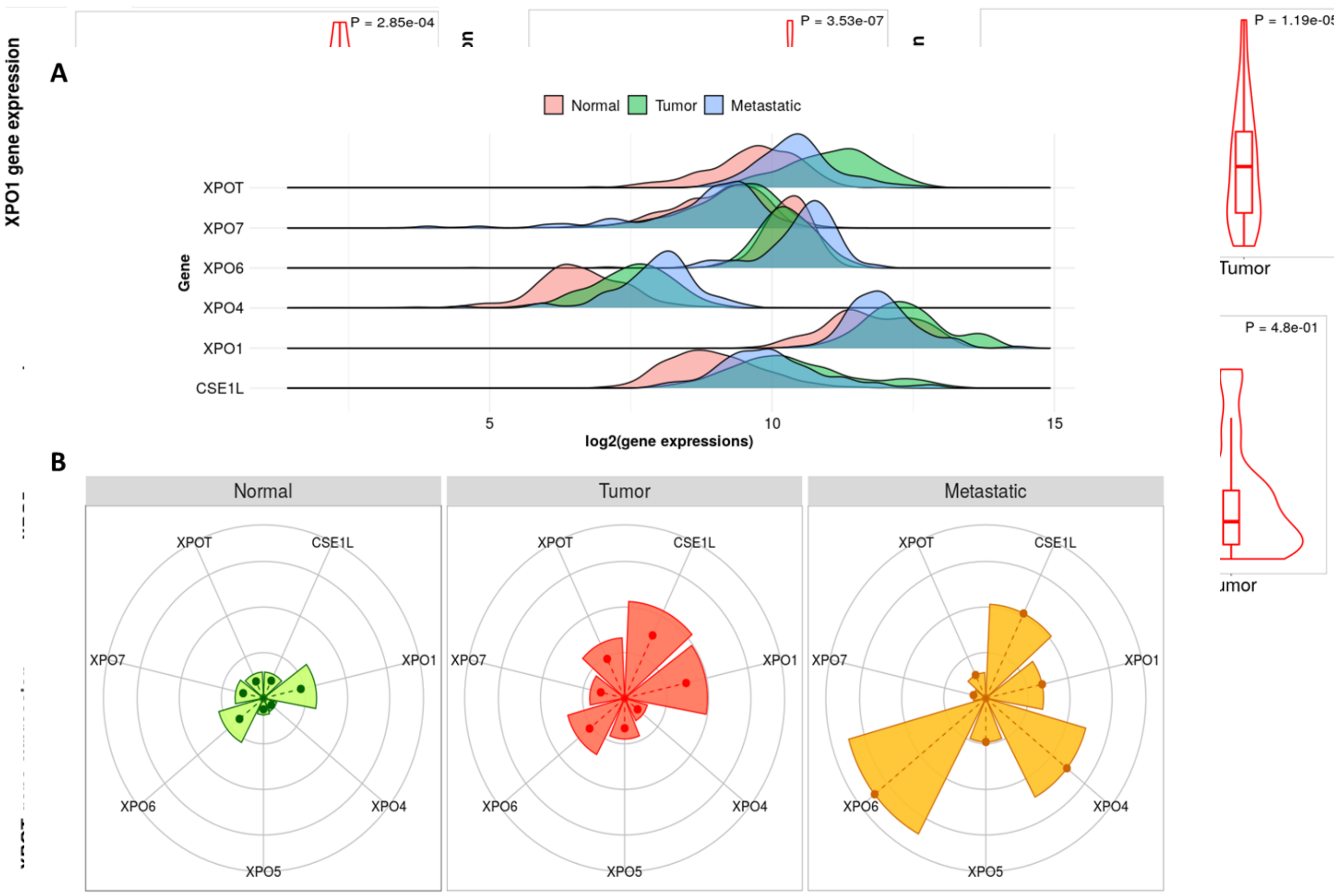


**Supplementary figure1:** Exportins expression between normal and COAD patients analysed by TNMplot.

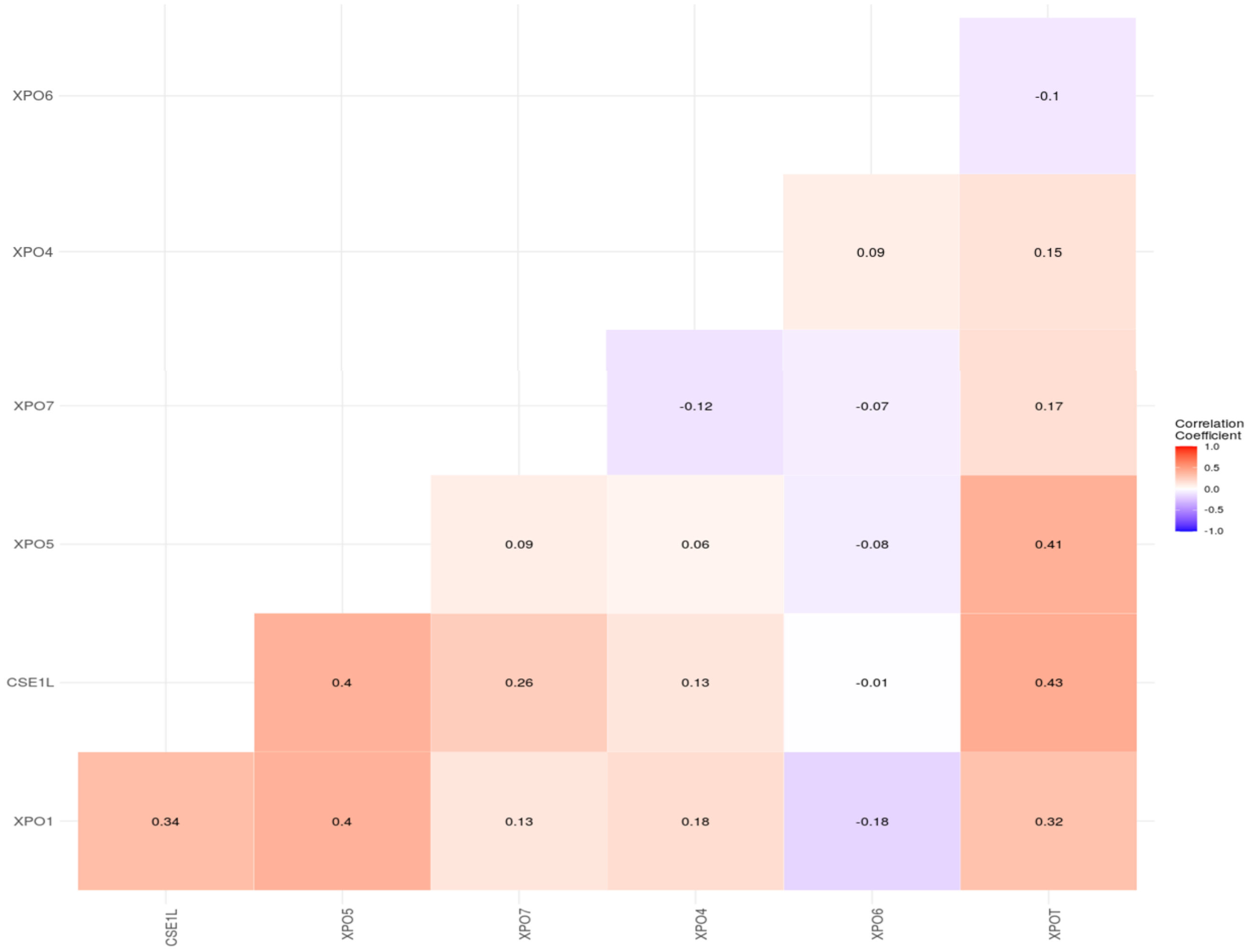


**Supplementary figure 2: A.** Exportins expression comparison between normal, tumor and metastatic sample of colon adenocarcinoma **B.** The target gram provide expression of exportins analysed by TNMplot. The size of segment represents mean value, length of dashed lines represent the median value.

**Supplementary figure 3:** Heatmap showing co-expression of exportins with respect to each other.

| Overall survival | | | Relapse free survival | |
|---|---|---|---|---|
| | Number of sample with low expression compared to median expression value | Number of sample with high expression compared to median expression value | Number of sample with low expression compared to median expression value | Number of sample with high expression compared to median expression value |
| XPO1 | 138 | 94 | 50 | 114 |
| XPO2 | 135 | 97 | 40 | 124 |
| XPO4 | 173 | 59 | 90 | 74 |
| XPO5 | 100 | 132 | 48 | 116 |
| XPO6 | 143 | 89 | 98 | 66 |
| XPO7 | 173 | 59 | 69 | 95 |
| XPOT | 97 | 135 | 79 | 85 |
| Total | **959** | **665** | **474** | **674** |

**Supplementary Table 1:** Number of patients used for the survival analysis

| Correlated Gene | Cytoband | spearman's corelation | p- value | q value |
|---|---|---|---|---|
| XPO1 | | | | |
| ATAD5 | 17q11.2 | 0.655 | 3.61E-48 | 7.19E-44 |
| PMS1 | 2q32.2 | 0.632 | 4.84E-44 | 4.81E-40 |
| WDR43 | 2p23.2 | 0.618 | 1.18E-41 | 7.80E-38 |
| TOP2A | 17q21.2 | 0.597 | 2.61E-38 | 1.30E-34 |
| BUB1 | 2q13 | 0.595 | 5.29E-38 | 2.10E-34 |
| UBXN6 | 19p13 | -0.585 | 1.81E-36 | 5.32E-33 |
| SASS6 | 1p21.2 | 0.585 | 1.87E-36 | 5.32E-33 |
| XRCC2 | 7q36.1 | 0.584 | 2.29E-36 | 5.38E-33 |
| CIP2A | 3q13.13 | 0.584 | 2.45E-36 | 5.38E-33 |
| WDR75 | 2q32.2 | 0.584 | 2.71E-36 | 5.38E-33 |

| | | | | |
|---|---|---|---|---|
| CSE1L | | | | |
| DPM1 | 20q13.13 | 0.802 | 6.11E-87 | 1.22E-82 |
| EIF2S2 | 20q11.22 | 0.792 | 1.27E-83 | 1.27E-79 |
| VAPB | 20q13.32 | 0.791 | 4.41E-83 | 2.93E-79 |
| NELFCD | 20q13.32 | 0.79 | 1.19E-82 | 5.93E-79 |
| RAE1 | 20q13.31 | 0.785 | 3.62E-81 | 1.44E-77 |
| TPX2 | 20q11.21 | 0.783 | 1.74E-80 | 5.78E-77 |
| CDK5RAP1 | 20q11.21 | 0.781 | 9.44E-80 | 2.68E-76 |
| TTI1 | 20q11.23 | 0.779 | 6.28E-79 | 1.56E-75 |
| MOCS3 | 20q13.13 | 0.77 | 3.15E-76 | 6.97E-73 |
| UBE2V1 | 20q13.13 | 0.769 | 1.07E-75 | 2.13E-72 |
| XPO4 | | | | |
| RNF6 | 13q12.13 | 0.781 | 7.87E-80 | 1.57E-75 |
| AKAP11 | 13q14.11 | 0.776 | 6.02E-78 | 5.99E-74 |
| PDS5B | 13q13.1 | 0.763 | 5.31E-74 | 3.52E-70 |
| PROSER1 | 13q13.3 | 0.755 | 1.60E-71 | 7.97E-68 |
| NUP58 | 13q12.13 | 0.751 | 2.09E-70 | 8.31E-67 |
| CENPJ | 13q12.12-q12.13 | 0.747 | 3.14E-69 | 1.04E-65 |
| MPHOSPH8 | 13q12.11 | 0.742 | 4.51E-68 | 1.20E-64 |
| NAA16 | 13q14.11 | 0.742 | 4.82E-68 | 1.20E-64 |
| TPP2 | 13q33.1 | 0.74 | 2.12E-67 | 4.68E-64 |
| GPALPP1 | 13q14.12 | 0.733 | 1.12E-65 | 2.24E-62 |
| XPO5 | | | | |
| RPL7L1 | 6p21.1 | 0.636 | 1.06E-44 | 2.11E-40 |
| HSP90AB1 | 6p21.1 | 0.621 | 3.89E-42 | 3.87E-38 |
| CSE1L | 20q13.13 | 0.599 | 1.48E-38 | 9.79E-35 |
| MDC1 | 6p21.33 | 0.584 | 2.75E-36 | 1.37E-32 |
| ANAPC1 | 2q13 | 0.582 | 5.62E-36 | 1.96E-32 |
| SRPK1 | 6p21.31 | 0.582 | 5.91E-36 | 1.96E-32 |
| ANKS1A | 6p21.31 | 0.579 | 1.53E-35 | 4.35E-32 |
| ABCF1 | 6p21.33 | 0.574 | 6.29E-35 | 1.56E-31 |
| RBL1 | 20q11.23 | 0.57 | 3.13E-34 | 6.93E-31 |
| POLR1A | 2p11.2 | 0.567 | 6.98E-34 | 1.39E-30 |
| XPO6 | | | | |
| GTF3C1 | 16p12.1 | 0.621 | 4.68E-42 | 9.31E-38 |
| SETD1A | 16p11.2 | 0.603 | 3.42E-39 | 3.41E-35 |
| HCFC1 | Xq28 | 0.594 | 8.88E-38 | 5.89E-34 |
| POLR3E | 16p12.2 | 0.576 | 4.01E-35 | 1.99E-31 |
| RNF40 | 16p11.2 | 0.553 | 5.10E-32 | 2.03E-28 |
| USP31 | 16p12.2 | 0.545 | 5.68E-31 | 1.88E-27 |
| TBC1D24 | 16p13.3 | 0.543 | 1.06E-30 | 3.02E-27 |
| ATXN2L | 16p11.2 | 0.537 | 7.11E-30 | 1.77E-26 |
| TBC1D10B | 16p11.2 | 0.536 | 9.06E-30 | 2.00E-26 |
| FBXL19 | 16p11.2 | 0.534 | 1.62E-29 | 3.22E-26 |
| XPO7 | | | | |
| CCAR2 | 8p21.3 | 0.889 | 4.58E-131 | 9.12E-127 |

| | | | | |
|---|---|---|---|---|
| CHMP7 | 8p21.3 | 0.859 | 9.88E-113 | 9.83E-109 |
| PCM1 | 8p22 | 0.846 | 5.21E-106 | 3.45E-102 |
| CCDC25 | 8p21.1 | 0.826 | 1.35E-96 | 6.71E-93 |
| VPS37A | 8p22 | 0.823 | 3.49E-95 | 1.39E-91 |
| ELP3 | 8p21.1 | 0.818 | 3.04E-93 | 1.01E-89 |
| CNOT7 | 8p22 | 0.814 | 9.01E-92 | 2.56E-88 |
| INTS9 | 8p21.1 | 0.805 | 2.56E-88 | 6.01E-85 |
| INTS10 | 8p21.3 | 0.805 | 2.72E-88 | 6.01E-85 |
| ERI1 | 8p23.1 | 0.802 | 4.04E-87 | 8.04E-84 |
| XPOT | | | | |
| MARS1 | 12q13.3 | 0.712 | 2.74E-60 | 5.46E-56 |
| NAA25 | 12q24.13 | 0.614 | 5.48E-41 | 5.45E-37 |
| NUP107 | 12q15 | 0.596 | 4.20E-38 | 2.78E-34 |
| PTPN11 | 12q24.13 | 0.588 | 6.08E-37 | 3.02E-33 |
| PSAT1 | 9q21.2 | 0.578 | 1.80E-35 | 7.15E-32 |
| MTHFD1L | 6q25.1 | 0.57 | 2.66E-34 | 8.84E-31 |
| DENR | 12q24.31 | 0.556 | 1.96E-32 | 5.58E-29 |
| PNPT1 | 2p16.1 | 0.552 | 8.68E-32 | 2.16E-28 |
| SLC7A5 | 16q24.2 | 0.551 | 1.08E-31 | 2.38E-28 |
| IARS1 | 9q22.31 | 0.55 | 1.24E-31 | 2.47E-28 |

**Supplementary Table 2:** Top 10 highly co-expressed gene with each exportin their spearman's rank correlation coefficient

| **Category** | **Term** | **Count** | **PValue** | **Genes** | **Fold Enrichment** |
|---|---|---|---|---|---|
| GOTERM_BP_DIRECT | GO:0060236~regulation of mitotic spindle organization | 4 | 5.12E-05 | TPX2, CENPJ, SASS6, RAE1 | 53.93838 |
| GOTERM_BP_DIRECT | GO:0034243~regulation of transcription elongation from RNA polymerase II promoter | 3 | 0.0024 | INTS10, INTS9, WDR43 | 40.45378 |
| GOTERM_BP_DIRECT | GO:0045070~positive regulation of viral genome replication | 3 | 0.004557 | CNOT7, VAPB, SRPK1 | 29.29412 |
| GOTERM_BP_DIRECT | GO:0007049~cell cycle | 6 | 0.007504 | MDC1, RBL1, SASS6, RAE1, CCAR2, HCFC1 | 4.840623 |
| GOTERM_BP_DIRECT | GO:0051301~cell division | 6 | 0.011031 | TPX2, CENPJ, PDS5B, RAE1, BUB1, ANAPC1 | 4.401707 |
| GOTERM_BP_DIRECT | GO:0006913~nucleocytoplasmic transport | 3 | 0.013626 | NUP107, RAE1, NUP58 | 16.65744 |
| GOTERM_BP_DIRECT | GO:2000003~positive regulation of DNA damage checkpoint | 2 | 0.013846 | CCAR2, TTI1 | 141.5882 |
| GOTERM_BP_DIRECT | GO:0046601~positive regulation of centriole replication | 2 | 0.017278 | CENPJ, SASS6 | 113.2706 |
| GOTERM_BP_DIRECT | GO:0006974~cellular response to DNA damage stimulus | 5 | 0.019673 | TOP2A, MDC1, SETD1A, CIP2A, CCAR2 | 4.783386 |
| GOTERM_BP_DIRECT | GO:0017196~N-terminal peptidyl-methionine acetylation | 2 | 0.020699 | NAA25, NAA16 | 94.39216 |

| | | | | | |
|---|---|---|---|---|---|
| GOTERM_BP_DIRECT | GO:1905832~positive regulation of spindle assembly | 2 | 0.027504 | CENPJ, SASS6 | 70.79412 |
| GOTERM_BP_DIRECT | GO:0006338~chromatin remodeling | 5 | 0.029284 | FBXL19, SETD1A, BUB1, TTI1, HCFC1 | 4.226514 |
| GOTERM_BP_DIRECT | GO:0001731~formation of translation preinitiation complex | 2 | 0.030889 | DENR, EIF2S2 | 62.9281 |
| GOTERM_BP_DIRECT | GO:2000234~positive regulation of rRNA processing | 2 | 0.034262 | WDR75, WDR43 | 56.63529 |
| GOTERM_BP_DIRECT | GO:0061014~positive regulation of mRNA catabolic process | 2 | 0.040975 | PNPT1, CNOT7 | 47.19608 |
| GOTERM_BP_DIRECT | GO:0042797~tRNA transcription from RNA polymerase III promoter | 2 | 0.040975 | GTF3C1, POLR3E | 47.19608 |
| GOTERM_BP_DIRECT | GO:0007059~chromosome segregation | 3 | 0.044112 | TOP2A, BUB1, SRPK1 | 8.849265 |
| GOTERM_BP_DIRECT | GO:0009303~rRNA transcription | 2 | 0.044314 | GTF3C1, MARS1 | 43.56561 |
| GOTERM_BP_DIRECT | GO:0050821~protein stabilization | 4 | 0.04563 | HSP90AB1, NAA16, TTI1, HCFC1 | 4.946314 |
| GOTERM_BP_DIRECT | GO:0016180~snRNA processing | 2 | 0.050957 | INTS10, INTS9 | 37.75686 |
| GOTERM_BP_DIRECT | GO:0002098~tRNA wobble uridine modification | 2 | 0.054262 | MOCS3, ELP3 | 35.39706 |
| GOTERM_BP_DIRECT | GO:0043254~regulation of protein complex assembly | 2 | 0.073854 | PTPN11, HCFC1 | 25.74332 |
| GOTERM_BP_DIRECT | GO:0007099~centriole replication | 2 | 0.07708 | CENPJ, SASS6 | 24.62404 |
| GOTERM_BP_DIRECT | GO:0060255~regulation of macromolecule metabolic process | 2 | 0.080295 | TOP2A, HSP90AB1 | 23.59804 |
| GOTERM_BP_DIRECT | GO:0007020~microtubule nucleation | 2 | 0.080295 | TPX2, CENPJ | 23.59804 |
| GOTERM_BP_DIRECT | GO:0006418~tRNA aminoacylation for protein translation | 2 | 0.080295 | MARS1, IARS1 | 23.59804 |
| GOTERM_BP_DIRECT | GO:0039702~viral budding via host ESCRT complex | 2 | 0.080295 | VPS37A, CHMP7 | 23.59804 |
| GOTERM_BP_DIRECT | GO:0051171~regulation of nitrogen compound metabolic process | 2 | 0.086692 | TOP2A, HSP90AB1 | 21.78281 |
| GOTERM_BP_DIRECT | GO:0045943~positive regulation of transcription from RNA polymerase I promoter | 2 | 0.086692 | WDR75, WDR43 | 21.78281 |
| GOTERM_MF_DIRECT | GO:0005515~protein binding | 62 | 4.70E-06 | TOP2A, MDC1, NUP107, MOCS3, HSP90AB1, CSE1L, INTS10, CCAR2, WDR43, PCM1, AKAP11, DENR, NELFCD, ANKS1A, UBXN6, TBC1D10B, PROSER1, VPS37A, CIP2A, RNF40, TTI1, HCFC1, ATXN2L, DPM1, SLC7A5, RBL1, POLR1A, RPL7L1, TBC1D24, UBE2V1, CHMP7, NUP58, GTF3C1, CCDC25, RNF6, PDS5B, ERI1, NAA25, IARS1, RAE1, BUB1, PMS1, ABCF1, PNPT1, ATAD5, FBXL19, XRCC2, SETD1A, PTPN11, ELP3, EIF2S2, SRPK1, TPX2, CNOT7, VAPB, PSAT1, CENPJ, SASS6, TPP2, NAA16, INTS9, MPHOSPH8 | 1.354911 |
| GOTERM_MF_DIRECT | GO:0003723~RNA binding | 15 | 5.68E-04 | TOP2A, PNPT1, HSP90AB1, SETD1A, WDR75, EIF2S2, | 2.823978 |

| | | | | | |
|---|---|---|---|---|---|
| | | | | CCAR2, WDR43, SRPK1, ATXN2L, CNOT7, RPL7L1, NELFCD, RAE1, ABCF1 | |
| GOTERM_MF_DIRECT | GO:0000175~3'-5'-exoribonuclease activity | 3 | 0.004733 | PNPT1, CNOT7, ERI1 | 28.72667 |
| GOTERM_MF_DIRECT | GO:0045296~cadherin binding | 6 | 0.00555 | ATXN2L, HSP90AB1, VAPB, PTPN11, CIP2A, HCFC1 | 5.20671 |
| GOTERM_MF_DIRECT | GO:0005524~ATP binding | 12 | 0.018971 | TOP2A, MOCS3, HSP90AB1, ATAD5, MTHFD1L, XRCC2, MARS1, IARS1, BUB1, PMS1, ABCF1, SRPK1 | 2.163827 |
| GOTERM_MF_DIRECT | GO:0043022~ribosome binding | 3 | 0.029146 | ERI1, NAA16, ABCF1 | 11.10765 |
| GOTERM_MF_DIRECT | GO:0060090~binding, bridging | 4 | 0.030835 | TPX2, PCM1, PTPN11, ANAPC1 | 5.785233 |
| GOTERM_MF_DIRECT | GO:0000049~tRNA binding | 3 | 0.033573 | MARS1, ELP3, IARS1 | 10.28486 |
| GOTERM_MF_DIRECT | GO:0003677~DNA binding | 10 | 0.048216 | TOP2A, GTF3C1, ATAD5, POLR1A, FBXL19, XRCC2, CCDC25, RNF6, PDS5B, PMS1 | 2.056972 |
| GOTERM_MF_DIRECT | GO:0016407~acetyltransferase activity | 2 | 0.051939 | ELP3, NAA16 | 37.02549 |
| GOTERM_MF_DIRECT | GO:0008135~translation factor activity, RNA binding | 2 | 0.075259 | EIF2S2, ABCF1 | 25.24465 |
| GOTERM_MF_DIRECT | GO:0017056~structural constituent of nuclear pore | 2 | 0.08833 | NUP107, NUP58 | 21.36086 |
| GOTERM_MF_DIRECT | GO:0019901~protein kinase binding | 5 | 0.092372 | TPX2, HSP90AB1, CENPJ, CDK5RAP1, PTPN11 | 2.87465 |
| GOTERM_CC_DIRECT | GO:0017101~aminoacyl-tRNA synthetase multienzyme complex | 2 | 0.034823 | MARS1, IARS1 | 55.68792 |
| GOTERM_CC_DIRECT | GO:0005694~chromosome | 4 | 0.049927 | MDC1, POLR1A, CIP2A, PDS5B | 4.76706 |
| GOTERM_CC_DIRECT | GO:0031965~nuclear membrane | 4 | 0.05184 | PCM1, NUP107, RNF6, NUP58 | 4.694001 |
| GOTERM_CC_DIRECT | GO:0048188~Set1C/COMPASS complex | 2 | 0.053312 | SETD1A, HCFC1 | 36.03336 |
| GOTERM_CC_DIRECT | GO:0032039~integrator complex | 2 | 0.062424 | INTS10, INTS9 | 30.62836 |
| GOTERM_CC_DIRECT | GO:0035097~histone methyltransferase complex | 2 | 0.077421 | SETD1A, HCFC1 | 24.50269 |
| GOTERM_CC_DIRECT | GO:0000776~kinetochore | 3 | 0.0899 | NUP107, CHMP7, BUB1 | 5.890069 |
| GOTERM_CC_DIRECT | GO:0042788~polysomal ribosome | 2 | 0.095105 | PNPT1, ABCF1 | 19.76023 |

**Supplementary table 3:** Various biological processes were identified involving top two co-expressed genes for each exportin using DAVID database. Top two co-expressed genes for each exportins are highlighted in different colors.

| GO terms | TERMS | Count | Subgroup | log 10 p-value | P-Value | Enrichment |
|---|---|---|---|---|---|---|
| GO:0051301 | cell division | 7 | biological processes | 2.55603562 | 0.002779485 | 4.91833905 |
| GO:007049 | cell cycle | 6 | biological processes | 2.045822697 | 0.008998649 | 4.636090045 |
| GO:0006974 | cellular response to DNA damage stimulus | 5 | biological processes | 1.643679211 | 0.022715421 | 4.581271412 |
| GO:0006338 | chromatin remodeling | 5 | biological processes | 1.472904365 | 0.033658568 | 4.047929367 |
| GO:0060236 | regulation of mitotic spindle organization | 4 | biological processes | 4.233729337 | 5.84E-05 | 51.65928907 |
| GO:0034243 | regulation of transcription elongation from RNA polymerase II promoter | 3 | biological processes | 2.582349299 | 0.002616078 | 38.7444668 |
| GO:0045070 | positive regulation of viral genome replication | 3 | biological processes | 2.30418491 | 0.004963809 | 28.05633803 |
| GO:0006913 | nucleocytoplasmic transport | 3 | biological processes | 1.829503636 | 0.014807999 | 15.95360398 |
| GO:0007059 | chromosome segregation | 3 | biological processes | 1.32125663 | 0.047724718 | 8.475352113 |
| GO:0046601 | positive regulation of DNA damage checkpoint | 2 | biological processes | 1.839743253 | 0.014462945 | 135.6056338 |
| GO:0005829 | cytosol | 42 | cellular component | 7.831332419 | 1.47E-08 | 2.223695142 |
| GO:0005737 | cytoplasm | 38 | cellular component | 5.408591011 | 3.90E-06 | 1.976574065 |
| GO:0005654 | nucleoplasm | 37 | cellular component | 8.964979509 | 1.08E-09 | 2.730819307 |
| GO:0005634 | nucleus | 34 | cellular component | 3.048544387 | 8.94E-04 | 1.655429805 |
| GO:0005730 | nucleolus | 10 | cellular component | 1.369710459 | 0.042686401 | 2.107527986 |
| GO:0005813 | centrosome | 7 | cellular component | 1.656008582 | 0.022079611 | 3.152227343 |
| GO:0032991 | macromolecular complex | 7 | cellular component | 1.542372686 | 0.028683181 | 2.965462428 |
| GO:0005635 | nuclear envelope | 6 | cellular component | 3.169428623 | 6.77E-04 | 8.456456044 |
| GO:0005643 | nuclear pore | 5 | cellular component | 3.477180929 | 3.33E-04 | 14.95699708 |
| GO:0031965 | nuclear membrane | 5 | cellular component | 1.935996736 | 0.011587861 | 5.616037219 |
| GO:0005515 | protein binding | 64 | molecular function | 5.596438518 | 2.53E-06 | 1.358657208 |
| GO:0003723 | RNA binding | 16 | molecular function | 3.63546664 | 2.31E-04 | 2.926179177 |
| GO:0005524 | ATP binding | 12 | molecular function | 1.634054114 | 0.023224474 | 2.102003711 |
| GO:0045296 | cadherin binding | 6 | molecular function | 2.201764528 | 0.00628399 | 5.057946429 |
| GO:0060090 | binding, bridging | 4 | molecular function | 1.478390662 | 0.033236045 | 5.619940476 |
| GO:0000175 | 3'-5'-exoribonuclease activity | 3 | molecular function | 2.29991444 | 0.00501286 | 27.90591133 |

| | | | | | | |
|---|---|---|---|---|---|---|
| GO:0043022 | ribosome binding | 3 | molecular function | 1.511880508 | 0.030769433 | 10.79028571 |
| GO:0000049 | tRNA binding | 3 | molecular function | 1.450647143 | 0.035428508 | 9.991005291 |
| hsa03013 | Nucleocytoplasmic transport | 9 | KEGG pathway | 7.515123821 | 3.05E-08 | 17.18650794 |
| hsa04110 | Cell cycle | 4 | KEGG pathway | 1.423710117 | 0.037695533 | 5.254473764 |
| hsa04914 | Progesterone-mediated oocyte maturation | 3 | KEGG pathway | 1.077081731 | 0.083737168 | 6.065826331 |
| hsa05014 | Amyotrophic lateral sclerosis | 5 | KEGG pathway | 1.036427803 | 0.091954333 | 2.832940869 |

**Supplementary table 4:** Gene Ontology and KEGG Pathway data
